# Activation of mu-opioid receptors slows pacemaking in hypothalamic A11 dopamine neurons

**DOI:** 10.64898/2026.07.08.737263

**Authors:** Angela F. Smith, Hannah N. Rust, Kathleen A. Sluka, Stephanie C. Gantz

## Abstract

Hypothalamic A11 dopamine neurons provide the only known source of spinal dopamine and critically modulate pain and motor systems. Yet, the electrophysiological properties of A11 neurons were unknown. Here, we characterized A11 dopamine neurons in mice using brain slice immunohistochemistry, and fluorescence-guided whole-cell patch-clamp and cell-attached electrophysiology. A11 dopamine neurons contained the enzymes necessary to synthesize dopamine, projected to the spinal cord, and were small, morphologically simple, and high resistance. Additionally, they received excitatory glutamatergic and inhibitory GABAergic synaptic input. Most A11 dopamine neurons fired action potentials spontaneously in a rhythmic ‘pacemaker’ manner at ∼5 Hz, while the remainder were quiescent at rest, but fired readily with somatic current injection. Pacemaking A11 dopamine neurons were differentiated from quiescent neurons by a net inward current at subthreshold potentials. Activation of mu-opioid receptors reduced the net inward current at subthreshold potentials via activation of potassium current but also decreased GABAergic synaptic currents onto A11 dopamine neurons. Using cell-attached recording to preserve the natural chloride gradient, we found mu-opioid receptor agonism reduced spontaneous action potential firing of A11 dopamine neurons. The results lay the necessary framework for future studies investigating synaptic and ion channel mechanisms underlying the excitability in A11 dopamine neurons in physiological and pathological conditions.

## Introduction

Dopamine is a highly evolutionarily conserved neurotransmitter, found in nearly every animal on Earth. From the ancient lamprey to human, the overall anatomical organization of the dopamine brain nuclei and their primary functions in regulating motor behavior and action selection are largely conserved. Early anatomical studies using rodent models defined nine distinct dopamine brain nuclei based on location and morphological characteristics, which were named A8 through A16 [1,2]. The focus of this study is the hypothalamic A11 dopamine neurons. Axon terminal projections of the A11 dopamine neurons are the only known source of dopamine release within the spinal dorsal horn [3–5] which crucially regulates nociceptive signaling (pain and itch) [3,6–10] and motor activity [11–13], with some circadian rhythmicity [10,14]. In animal models, dysfunction in A11 dopamine neurons and spinal dopamine signaling have been implicated in chronic pain, hyperalgesia and allodynia, migraine, and restless legs syndrome [14–22]. Despite these central functions, the electrophysiological and firing properties of A11 dopamine neurons have not been described.

Here, using fluorescence-guided whole-cell patch-clamp electrophysiological recordings in acute mouse brain slices, we show that A11 dopamine neurons are morphologically small and simple, with high membrane resistance. Most A11 dopamine neurons fired action potentials spontaneously in a slow, rhythmic ‘pacemaker’ manner of ∼5 Hz. A subset of A11 dopamine neurons were silent at rest but fired easily with somatic current injection. The subthreshold drive to fire action potentials in the pacemaking neurons was provided by a net inward current that was reduced by activation of mu-opioid receptors.

## Materials and Methods

### Animals

Male and female C57BL/6J mice, (stock #000664), TH-Cre-1 mice (#008601) [23], and TH-2A-CreER mice (#025614) [24] (The Jackson Laboratory) and TH-IRES-Cre mice (EM:00254) [25] were group housed on a 12/12 light/dark cycle and used during the light cycle. Mice were crossed with Ai9 or Ai14 Cre reporter mice (Jackson Laboratory, #007909, #007914) [26]. 8-week-old TH-2A-CreER::Ai14 mice received four daily injections of tamoxifen (75 mg/kg, i.p, 2 days on, one day off) to induce Cre recombination and recovered for >3 weeks before use. All experiments were conducted with the approval of the protocols (K.A.S and S.C.G) to the University of Iowa Institutional Animal Care and Use Committee.

### Brain slice preparation and electrophysiological recordings

Whole-cell patch-clamp recordings were made from tdTomato^+^ A11 dopamine neurons from TH-IRES-Cre::Ai14/Ai9 and TH-2A-CreER::Ai14 mice. Mice were deeply anesthetized with isoflurane and euthanized via decapitation. Brains were removed rapidly and placed in warm, bubbled (95/5% O_2_/CO_2_) modified Krebs’ buffer containing (in mM), NaCl (126), KCl (2.5), MgCl_2_ (1.2), CaCl_2_ (1.2), NaH_2_PO_4_ (1.2), NaHCO_3_ (21.5), and D-glucose (11), containing 5 µM MK-801 to increase slice viability. Coronal slices (220 µm) containing A11 (plates 49-52 [27]) were made, stored at 28 °C in the same solution, and recovered for >30 min.

Electrophysiological recordings were made in modified Krebs’ buffer. For all recordings except those of GABA_A_ receptor-mediated IPSCs, the internal solution contained (in mM), K-methyl sulfate (124.56), KCl (7), NaCl (5.3), K-HEPES (7.07), MgCl_2_ (0.0215), CaCl_2_ (0.2245), EGTA (0.45), Na-GTP (0.26), Na-ATP (4.87), Na-creatine phosphate (4.59), and 0.1% Biotin ethylenediamine [28,29], pH 7.25 with KOH, mOsm ∼278. After recording, the brain slices were paraformaldehyde-fixed and immunostained for TH. Reported voltages are corrected for a liquid junction potential of −8 mV between the internal and external solutions. For complete methodology, see supplemental materials and methods.

### Retrograde tracing

Retrograde tracing was performed as previously described [30]. In brief, C57BL6/J mice were anesthetized with isoflurane, a laminectomy was performed, and 4% Fluoro-Gold (Fluorochrome) was applied to the dorsal spinal cord at the L4-L6 level. After 3 weeks, mice were euthanized, and tissue was processed for immunohistochemistry. For complete methodology, see supplemental materials and methods.

### Immunohistochemistry and confocal microscopy

Mice were euthanized for transcardial perfusion. Brains were extracted, post-fixed, and sectioned for immunohistochemistry. Slices were immunostained for TH, AADC, TpH2, DβH, and/or Fluoro-Gold. Slices used for electrophysiological recordings were placed in 4% PFA for 1 h following recordings and immunostained for TH and Neurobiotin. For complete methodology, see supplemental materials and methods.

Fluorescent images were obtained on a confocal microscope (Zeiss). Cell counts were made using FIJI (ImageJ) software. Specificity and penetrance were measured as the percentage of tdTomato^+^ neurons expressing TH and the percentage of TH^+^ neurons expressing tdTomato, respectively. Morphology measurements were calculated with the Simple Neurite Tracer plug-in for FIJI.

### Materials

See supplemental materials and methods.

### Experimental design and statistical analysis

See supplemental materials and methods.

## Results

### A11 dopamine neurons project to the spinal cord

A11 neurons were identified in coronal mouse brain slices by their location in the dorsal posterior hypothalamus, adjacent to the third ventricle and medial to the fasciculus retroflexes (Fig. 1A, B) [3,31,32]. A11 neurons were immunostained for the presence of tyrosine hydroxylase (TH, Fig. 1B), the rate-limiting enzyme in dopamine synthesis [4,32]. Consistent with prior reports [11,32], L-amino acid decarboxylase (AADC), which converts L-DOPA to dopamine, was found in 94% of TH^+^ A11 neurons (Fig. S1A-C). No neurons in A11 immunostained for dopamine-β-hydroxylase (DβH, required for noradrenaline synthesis, as confirmed by immunostaining in locus coeruleus, Fig. S1D, E)[1,33,34] or tryptophan hydroxylase 2 (TpH2, required for serotonin synthesis, as confirmed by immunostaining in dorsal raphe nucleus, Fig. S1F, G)[1]. Taken together, these data indicate that A11 TH^+^ neurons contain the enzymes required for dopamine synthesis, but not noradrenaline or serotonin, and will henceforth be referred to as “A11 dopamine neurons”.

**Fig 1.**
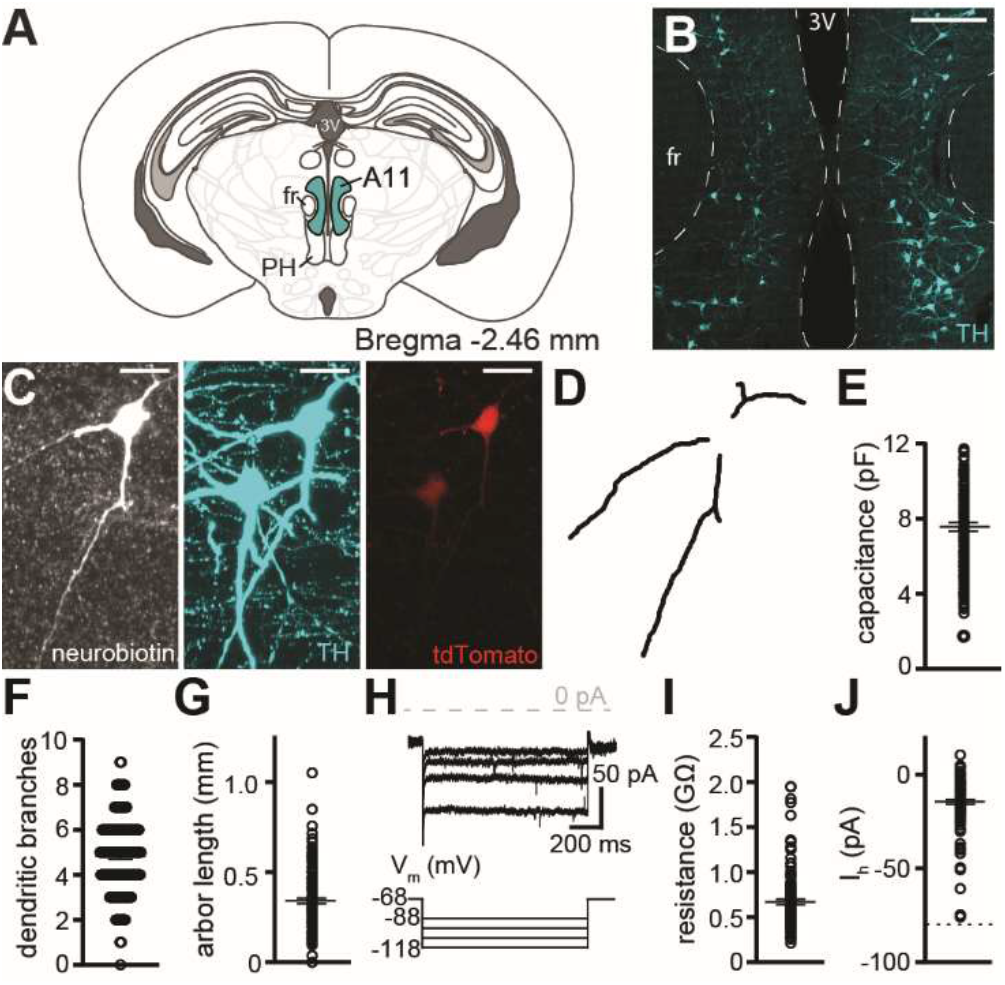
A11 dopamine neurons are small, morphologically simple, and high resistance. **A** Schematic of coronal mouse brain section containing A11 dopamine neurons. Adapted from Franklin and Paxinos (2008)[27]. Abbreviations: Third ventricle (3V), fasciculus retroflexes (fr), posterior hypothalamus (PH). **B** Maximum intensity projection confocal image of A11 dopamine neurons labeled by immunostaining for tyrosine hydroxylase (TH, cyan); Scale bar 200 μm. **C, D** Representative maximum intensity projection and reconstruction of an A11 dopamine neuron labeled with biotin (grey), TH (cyan), and tdTomato (red); scale bars 20 μm. **E** Capacitance of A11 dopamine neurons (n=144). **F** Total number of dendritic branches on A11 dopamine neurons (n=136). **G** Dendritic arbor length for A11 dopamine neurons (n=136). **H** Representative voltage-clamp trace showing the current induced by voltage steps to −88, −98, −108, and −118 mV from V_hold_ (−68 mV). **I** Input resistance measured by a −50 mV step from V_hold_ −68 mV (n=119). **J** Hyperpolarization-induced current (I_h_) from a voltage step to −118 mV from −68 mV. No neurons had I_h_ > −80 pA (dotted line) (n=119). Line and error bars represent means±SEM.

It is well-established that A11 dopamine neurons project to the spinal cord [4,5,31,32,35–38]. To determine the percentage of A11 dopamine neurons that project to the lumbosacral level of the spinal cord, Fluoro-Gold, a retrograde axonal tracer, was applied at the L4-L6 level of the spinal cord as previously described [30]. After allowing 21 days for transport, brain sections containing A11 were immunostained for TH and Fluoro-Gold. Fluoro-Gold was detected in 71% of A11 dopamine neurons (Fig. S2A, B). Thus, the majority of A11 dopamine neurons project to the lumbosacral spinal cord.

### Available Cre lines have variable specificity for A11 dopamine neurons

Since A11 is a small and sparse nucleus (Fig. 1B) [5,38], we utilized fluorescence-guided patch-clamp electrophysiology. Initially, TH-Cre-1, TH-IRES-CRE, and tamoxifen-inducible TH-2A-Cre mice were crossed with Ai14 Cre reporter mice that produce cytosolic tdTomato in Cre^+^ neurons. In A11 from TH-Cre-1::Ai14 and TH-IRES-Cre::Ai14 mice, only 30% and 40% of tdTomato^+^ neurons immunostained for TH, respectively (Fig. S3A-D), consistent with a prior report [11]. In A11 from the TH-2A-CreER::Ai14 mice, 67% of tdTomato^+^ neurons expressed TH (Fig. S3E, F). All three lines produced high penetrance of tdTomato expression in A11 dopamine neurons (Fig. S3B, D, F). Morphology and electrophysiology data below were collected from TH-IRES-Cre::Ai14/Ai9 (n=23 neurons) and TH-2A-CreER::Ai14 (n=137 neurons) mice.

### A11 dopamine neurons are small, morphologically simple, and high resistance

tdTomato^+^ A11 dopamine neurons in live brain slices were filled with 0.1% Neurobiotin [28,29], fixed, and immunostained for TH (Fig. 1C). The morphology of the dendritic arbor was reconstructed by tracing dendritic processes in *x*, *y*, and *z* dimensions (Fig. 1D). On average, the capacitance of the A11 dopamine neurons was ∼8 pF (Fig. 1E), indicating a small soma [5,32,38]. Given the limitations of somatic recording, capacitance measurements may not include distal dendrites. Therefore, the dendritic arbor was measured from the 3-dimensional digital reconstruction. A11 dopamine neurons had ∼5 dendritic branches, primarily originating from the soma and with only 1-2 higher order branches on average (Fig. 1D, F), and a total dendritic arbor length of ∼342 µm (Fig. 1G). The input resistance of A11 dopamine neurons was ∼670 MΩ when measured by a voltage-step from V_hold_ −68 mV to −118 mV (Fig. 1H, I). After the instantaneous change in current which reflects input resistance, there was little change in current in response to hyperpolarizing voltage-steps (Fig. 1H, J); unlike most midbrain dopamine neurons, which have a large hyperpolarization-activated inward (I_h_) current [39,40].

### A11 dopamine neurons can fire action potentials in a rhythmic pacemaker manner

Ventral midbrain dopamine neurons are known for their intrinsic excitability and rhythmic ‘pacemaker’-like action potential firing [41]. Using whole-cell current-clamp recording, we found that 79% of A11 dopamine neurons fired spontaneously at 5.5±0.3 Hz with a high pacemaker-like regularity (coefficient of variation: 0.36±0.04) (Fig. 2A, B). 18% of A11 dopamine neurons were either completely quiescent or fired a few action potentials before becoming silent (Fig. 2A, B). 3% appeared to be in depolarization block (n=4, Fig. 2B) with a resting membrane potential of −39.6±1.0 mV. The percentage of A11 dopamine neurons that were spontaneously active was greater in brain slices from male mice (92% vs. 61%, p<0.0001), but there were no differences in the frequency of firing or action potential shape (Table S1). Pacemaking was not a defining characteristic of A11 dopamine neurons, as 25% of TH^-^ neurons displayed pacemaker-like firing, while the majority were quiescent (Fig. S4). Using somatic current injection (−40 pA, 500 ms), spontaneously active pacemaker neurons were hyperpolarized. 64/87 neurons paused firing and were hyperpolarized to −77.3±1.8 mV. The remaining 23/87 neurons continued to fire, albeit more slowly, and were hyperpolarized to −63.7±1.2 mV (Fig. S5A). These data suggest that the spontaneous activity of pacemaker neurons can be stopped with sufficient hyperpolarization.

**Fig 2.**
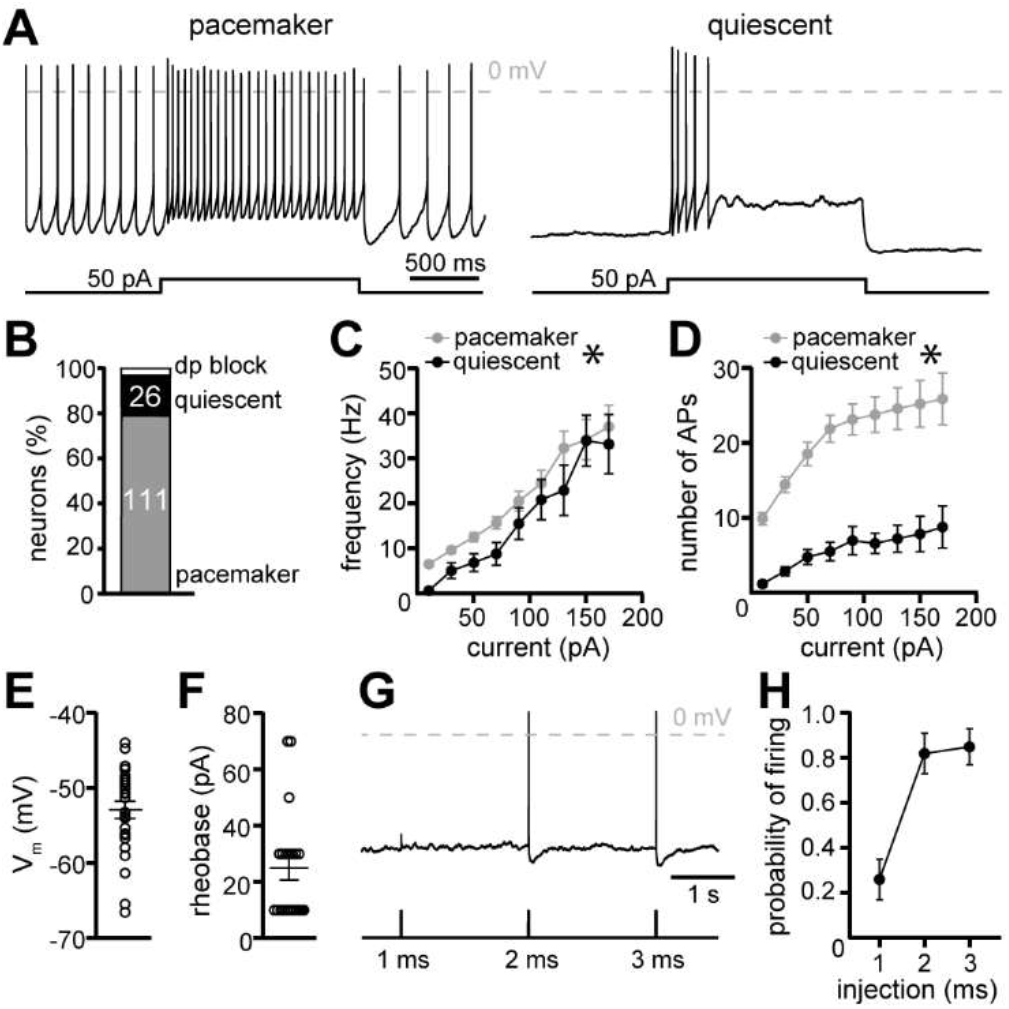
A11 dopamine neurons can fire action potentials in a rhythmic pacemaker manner. **A** Representative current-clamp traces of A11 dopamine neurons that fired action potentials spontaneously (‘pacemakers’) or were silent (‘quiescent’). Both types fired action potentials during somatic current injection (50 pA, 1.5 s). **B** Percentage of A11 dopamine neurons that fired spontaneously (pacemaker), were silent (quiescent), or in depolarization block (dp). Inset numbers represent n of neurons. n=141. **C** Frequency while firing during a somatic current injection was generally lower in quiescent neurons than pacemaker (p=0.0020, n=20-47, ordinary two-way ANOVA, main effect of type) but did not differ statistically at any one current injection increment (Sidak’s multiple comparison test). **D** Number of action potentials was significantly increased in pacemaker neurons. (p<0.0001, n=20-47, ordinary two-way ANOVA, main effect of type). **E** Resting membrane potential of quiescent A11 dopamine neurons (n=26). **F** Approximate rheobase of quiescent A11 dopamine neurons (n=20). **G** Representative trace of action potential firing in a quiescent neuron to brief current injections of increasing duration (150 pA, 1, 2, or 3 ms). **H** Probability of firing in quiescent neurons following 1, 2, and 3 ms of somatic current injection. (150 pA, n=13). Line and error bars represent means±SEM. * denotes statistical significance, ‘ns’ indicates not significant.

Using somatic current injection (1.5 s, 20 pA intervals) to evoke action potential firing, both pacemaker and quiescent neurons fired similar maximum frequencies, up to ∼40 Hz (Fig. 2C). However, quiescent neurons tended to fire slightly slower (Fig. 2C), and pacemaker neurons sustained firing for a longer duration, thereby firing more action potentials during current injection (Fig. 2A, D). Upon current injection, quiescent neurons fired <10 action potentials, then remained depolarized stably (Fig. 2A, D). The quiescent neurons were still readily excitable, with an average resting membrane potential of −53 mV (Fig. 2E) and an average rheobase of 25 pA (Fig. 2F). Further, quiescent neurons had a high probability of firing a single action potential in response to brief (2 ms) somatic current injection (150 pA, Fig. 2G, H). The threshold of the action potential from quiescent neurons was more depolarized than the threshold of the action potential from pacemaker neurons (Fig. S6A, B), but there were no differences in half-width, peak height, or the magnitude of the after-hyperpolarization (Fig. S6C-E).

### Subthreshold sodium inward current contributes to pacemaking in A11 dopamine neurons

Voltage-ramps from −50 mV to −130 mV (0.2 mV/ms) were used to record subthreshold current in pacemaker and quiescent neurons (Fig. 3A). At −50 mV, there was a net inward current of −63.0±5.1 pA in pacemaker neurons that was absent in quiescent neurons (−1.1±6.9 pA) (Fig. 3A, B). In pacemaker neurons, the magnitude of the net inward current was correlated positively to the spontaneous firing frequency (Fig. 3C).

**Fig 3.**
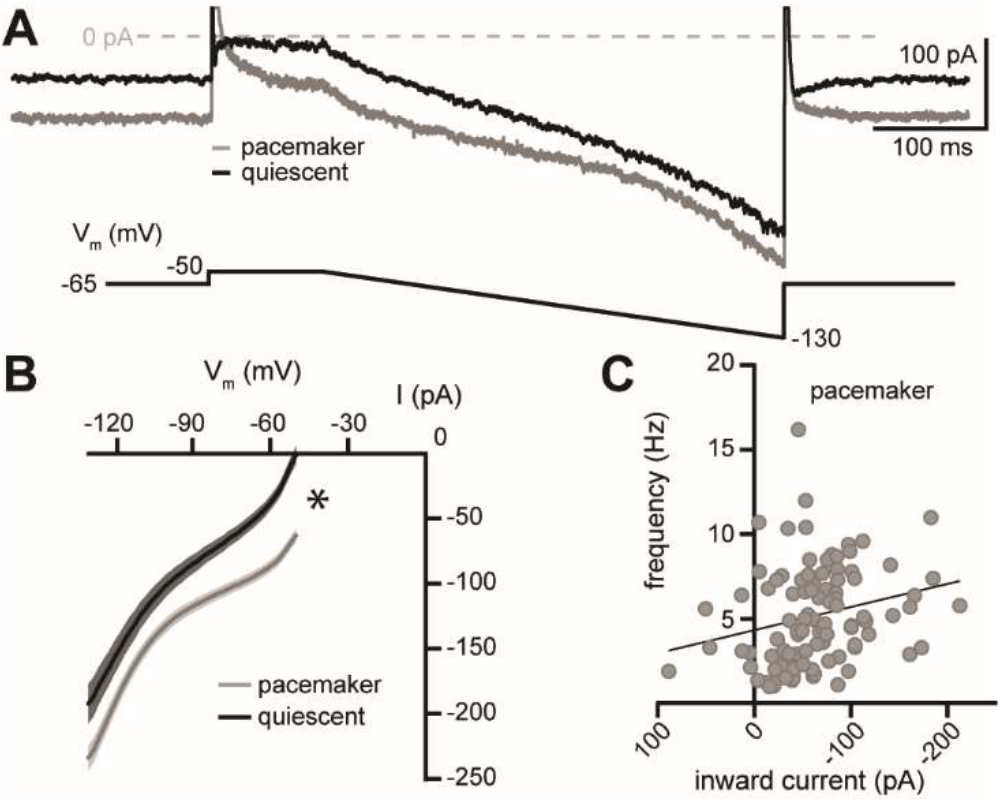
Pacemaker A11 dopamine neurons have a net inward current at subthreshold potentials. **A** Representative voltage-clamp traces of current induced by a voltage-ramp from −50 mV to −130 mV (0.2 mV/ms) in a pacemaker (grey) and quiescent (black) neuron. **B** Current-voltage plot (2 mV bins) generated by a voltage-ramp from −50 mV to −130 mV (0.2 mV/ms) showing a net inward current at between −50 and −56 mV that was present in pacemaker but not quiescent neurons (ordinary two-way ANOVA, main effect of type: p<0.0001, Sidak post-hoc tests: −50 to −70 mV, p<0.05, n=24-99). **C** Frequency of spontaneous firing in pacemaker neurons was correlated to the magnitude of the net inward current at −50 mV (simple linear regression, R^2^=0.06, p=0.0191, n=98). Line and error bars represent means±SEM. * denotes statistical significance.

Currents from both pacemaker and quiescent neurons showed inward rectification at potentials more negative to the reversal potential for potassium (Fig. 3B, calculated E_K_: −106 mV), likely due to “leak” potassium channels. To determine whether subthreshold inward current was minimal in quiescent neurons due to a competing outward potassium current, extracellular potassium was increased from 2.5 mM to 10.5 mM, which reduces the chemical driving force of potassium (calculated E_K_: −68 mV). At V_hold_ −65 mV, there was an apparent inward current due to a reduction in leak potassium current in quiescent neurons (∼18 pA), but not pacemaker neurons (Fig. S7). To determine whether subthreshold inward current in pacemaker neurons was due to sodium entry, 77% of extracellular sodium was replaced by N-methyl D-glucamine (NMDG, 115 mM). Application of NMDG-containing extracellular solution caused an apparent outward current in pacemaker neurons due to a reduction in leak sodium current (∼-60 pA, Fig. S8). The current-voltage relationship of the NMDG-sensitive current was largely voltage-independent (Fig. S8). Thus, sodium is the prominent charge carrier of subthreshold inward current in pacemaking A11 dopamine neurons.

### Activation of mu-opioid receptors reduces subthreshold inward current in A11 dopamine neurons

Despite the prominent role of A11 dopamine neurons in nociception [3,21], there are no reports of direct action of opioids on A11 dopamine neurons. Using the same voltage ramps, we measured subthreshold current before and during application of a mu-opioid receptor agonist, DAMGO (10 µM). In 4/12 neurons, DAMGO had no effect on subthreshold current (Fig. 4A). In 8/12 neurons, at V_hold_ −50 mV, DAMGO produced an apparent outward current (Fig. 4B-C). Over the full voltage range tested there was only a subtle increase in conductance to DAMGO (Fig. 4D), which may be explained by a simultaneous increase in one conductance (e.g. K^+^ conductance) and a decrease in another (e.g. Na^+^ conductance) [42]. On average, the DAMGO-induced outward current was quite small at potentials more negative to V_hold_ −100 mV and did not reverse polarity (Fig. 4F), suggesting that activation of mu-opioid receptors likely affects conductance carried by more than one ionic species. Replacing extracellular sodium with NMDG had no effect on the outward current induced by DAMGO (7/9 responded, Fig. 4E), suggesting that DAMGO had little effect on sodium conductance (Fig. S9). In 10.5 mM extracellular potassium, the DAMGO-induced outward current was comparable to the outward current in control conditions (8/11 responded, Fig. 4E) yet reversed polarity at −80 mV and was inward at more negative potentials (Fig. 4F) These data suggest that activation of potassium channels is a primary target of mu-opioid receptors. However, since the current reversed polarity at a potential more negative than E_K_ (−68 mV), there may also be minor involvement of another unidentified conductance.

**Fig 4.**
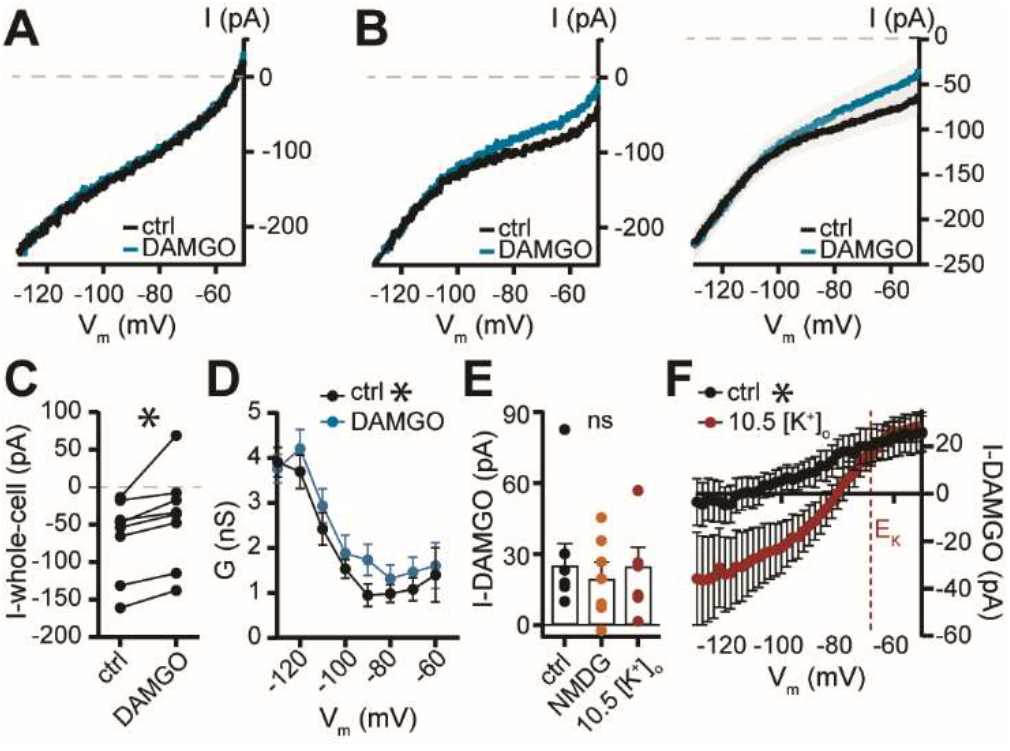
Activation of mu-opioid receptors reduces the net inward current at subthreshold potentials via activation of potassium current. **A,B** Current-voltage plots generated by a voltage-ramp from −50 mV to −130 mV (0.2 mV/ms) in control conditions (ctrl) and after application of an mu-opioid receptor agonist, DAMGO (10 μM) in (**A**) a neuron that had no response and (**B**) in a neuron in which DAMGO reduced the net inward current (left) also shown in grouped data (responders, right). **C** In neurons that responded to DAMGO, there was a significant decrease in the net inward current at V_hold_ −50 mV (Wilcoxon signed rank test, p=0.0078, n=8). **D** Conductance-voltage plot (10 mV bins) in control conditions (ctrl) and after application of DAMGO in responders (two-way repeated measures ANOVA, main effect of drug: p=0.0005, Sidak post-hoc tests: not significant. n=8). **E** Magnitude of the DAMGO-induced current (I-DAMGO) at V_hold_ −50 mV in control conditions (n=8), NMDG (n=7), and 10.5 mM extracellular potassium (n=8) (Kruskal-Wallis test, p=0.8958). **F** Current-voltage plots of the DAMGO-induced current (I-DAMGO) in control conditions (n=8, 2.5 mM extracellular potassium) and in 10.5 mM extracellular potassium (n=8, Ordinary two-way ANOVA, main effect of potassium concentration, p<0.0001). Red dashed line represents E_K_ in 10.5 mM extracellular potassium. Line and error bars represent means±SEM. * denotes statistical significance, ‘ns’ indicates not significant.

### Mu-opioid receptor agonism inhibits GABAergic synaptic input onto A11 dopamine neurons

The posterior hypothalamus receives innervation from multiple cortical, thalamic, hypothalamic, and brainstem regions [43,44] and contains local GABA and glutamate neurons [45,46], but there is no prior electrophysiological evidence of synaptic input to A11 dopamine neurons. While recording in the absence of synaptic blockers, spontaneous synaptic currents were observed. In the presence of a GABA_A_ receptor blocker, picrotoxin, spontaneous glutamatergic excitatory postsynaptic currents (sEPSCs) were recorded at a frequency of ∼1 Hz. Application of an AMPA/Kainate receptor antagonist, NBQX (Fig. S10A, B), or an AMPA receptor-specific antagonist, GYKI-52466 (n=4) blocked the sEPSCs, confirming the sEPSCs were produced by activation of postsynaptic AMPA receptors. Two populations of synaptic events, ‘fast’ and ‘slow’ were noted (Fig. S10C). Fast sEPSCs had a shorter time-to-peak and an accelerated rate of current decay (τ-decay) than slow sEPSCs (Fig. S10D, E). In principle, slower synaptic events may arise from more distal synapses and subsequently slow and attenuate before reaching the soma due to passive cable properties [47,48]. Indeed, dendritic arbor length was positively correlated with greater occurrence of slow sEPSCs (Fig. S10F) and slow sEPSCs had a smaller amplitude than fast sEPSCs (Fig. S10G) [47]. Using a high-chloride-containing internal solution, in the presence of NBQX, spontaneous GABAergic inhibitory postsynaptic currents (sIPSCs) were recorded at a frequency of ∼1.5 Hz (Fig. S11A-C). Application of DAMGO (10 µM) caused a significant reduction in the frequency and amplitude of sIPSCs (Fig. S11A, C, D), suggesting that activation of presynaptic mu-opioid receptors inhibits GABA release onto A11 dopamine neurons.

### Activation of mu-opioid receptors slows pacemaking in A11 dopamine neurons

To determine the net effect of DAMGO on A11 dopamine neuron pacemaking, fluorescence-guided cell-attached recordings were made to record action potential firing while maintaining the natural chloride gradient underlying GABA_A_ receptor-mediated inhibition. Following recording, slices were fixed and immunostained for TH. Recorded neurons were identified by location and morphology (Fig. 5A). Only spontaneously active TH^+^ neurons were analyzed. In cell-attached mode, A11 dopamine neurons fired at ∼3 Hz (Fig. 5B, C) with high regularity (coefficient of variation: 0.39±0.1, n = 11). Application of DAMGO (10 µM) reduced the frequency of spontaneous action potential firing (Fig. 5B-D); an effect that desensitized within a few minutes of the continued presence of DAMGO.

**Fig 5.**
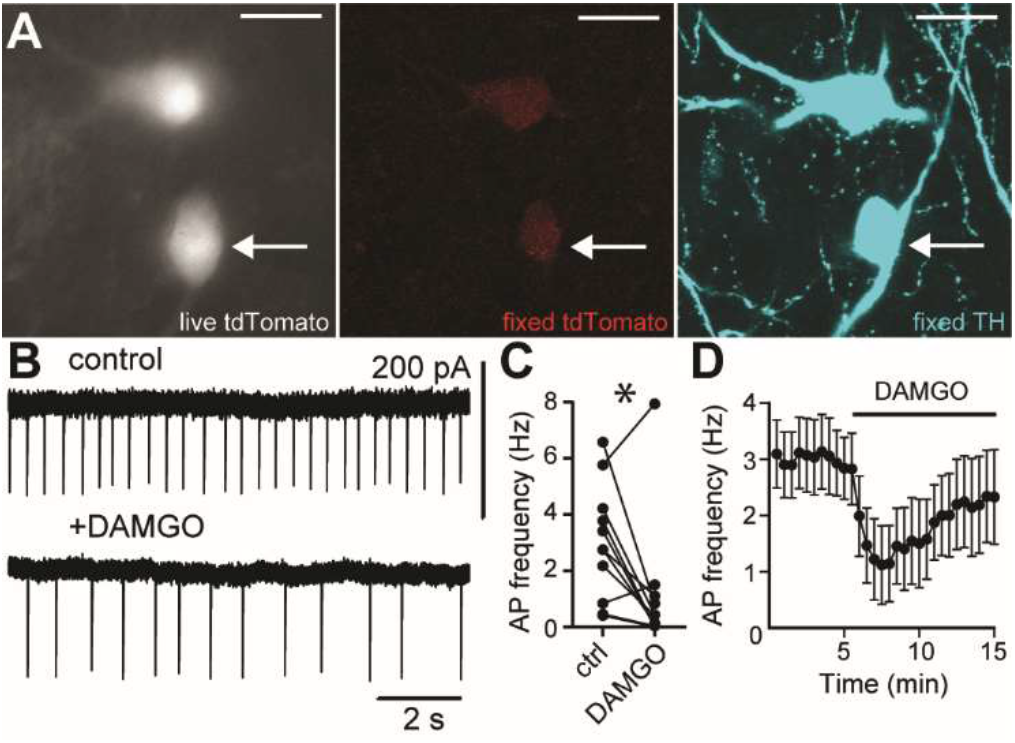
Activation of mu-opioid receptors slows pacemaking in A11 dopamine neurons. **A** Representative image of a tdTomato^+^ A11 neuron in a live brain slice (grey, arrow). Maximum intensity projection confocal images of the same A11 dopamine neuron following fixation, labeled with tdTomato (red) and tyrosine hydroxylase (TH, cyan); scale bar 20 µm. **B-D** DAMGO (10 μM) reduced the frequency of spontaneous action potential firing, shown in (**B**) a representative cell-attached recording of pacemaking in an A11 dopamine neuron, (**C**) grouped data (Wilcoxon matched-pairs signed rank test, p=0.0312, n=11 cells from 5 mice), and (**D**) an averaged time course. Line and error bars represent means±SEM. * denotes statistical significance.

## Discussion

### Distinguishing features of A11 dopamine neurons

A11 dopamine neurons are the only known source of dopamine to the spinal cord [3,4,36]. Their activity likely coordinates the release of spinal dopamine, where subsequent activation of dopamine receptors crucially regulates nociception [3,7,21,22] and motor activity [11–13]. Despite this critical role, the electrophysiological properties of A11 dopamine neurons had not been described. One major obstacle in recording the electrical activity of these neurons in acute mouse brain slices is that A11 is a small nucleus with less than two-hundred dopamine neurons [38]. For comparison, the noradrenergic LC, known for its small size, has over one-thousand neurons [49]. With the advent of site-specific recombinase systems, Cre-driven fluorescent labeling enables targeted whole-cell patch-clamp electrophysiological recordings. Yet still, the lack of specificity in Cre expression can limit experimental efficiency and interpretation [11,50]. For midbrain dopamine neurons, the use of a DAT-Cre transgenic mouse line greatly improves specificity over the TH-Cre lines [50]. However, the use of DAT-Cre mice may not be suitable for targeting A11 dopamine neurons. Multiple studies report that A11 dopamine neurons do not contain DAT [4,32,37,51,52], nor express Cre-dependent fluorescence when DAT-Cre mice are crossed with a Cre reporter line [11,53]. In the present study, we report that an inducible TH-Cre line (TH-2A-CreER) [24] nearly doubled the specificity of Cre expression for A11 dopamine neurons when compared to other available lines and may be the best Cre-driver line available currently to target A11 dopamine neurons.

Research in the last two decades has revealed marked heterogeneity within the midbrain dopamine neurons [5,41,54–56] and even greater heterogeneity amongst olfactory bulb and hypothalamic dopamine neurons in terms of morphology [40,57,58], terminal projections and synaptic innervation [57–61], electrophysiological profile [40,57,58,61–63], expression of calcium binding proteins [35,64–66] and expression of neuropeptides [4,60]. A11 dopamine neurons share some common features but are also likely heterogenous. Here, we report that A11 dopamine neurons were uniformly small with simple dendritic arbors. In A11, ∼94% of TH^+^ neurons also contained AADC, which is the final enzyme in the biosynthesis of dopamine. In humans, TH and AADC also colocalize in A11, suggesting this is a conserved trait [67]. Retrograde tracing studies suggest that the majority, if not all, of A11 dopamine neurons project to the spinal cord. Here we show that 71% of A11 dopamine neurons project to the lumbar and sacral segments. This is likely an underestimate, as only a portion of the lumbar spinal cord was exposed to Fluoro-Gold. Prior studies report that 30-70% of A11 dopamine neurons project to the C2, C5, L1, L4, or S1 segments [5,37,38]. Whether sub-populations of A11 dopamine neurons project to specific segments is unknown.

While the posterior hypothalamic region receives innervation from multiple cortical, thalamic, hypothalamic, and brainstem regions [43,44] and contains local GABA and glutamate neurons [11,45,46], synaptic connections onto A11 dopamine neurons have not been established. Here we report spontaneous glutamatergic EPSCs and GABAergic IPSCs recorded in A11 dopamine neurons at a low frequency of ∼1-1.5 Hz. The frequency of synaptic events is likely greater *in vivo*, due to some loss of synaptic input inherent to brain slice preparation. Future studies are required to determine the origin of glutamate synaptic input onto A11 dopamine neurons.

### Pacemaking in A11 dopamine neurons

Spontaneous action potential firing appears to be a common trait of dopamine neurons, although the pattern varies by region [40,57,58,63]. In brain slices, midbrain and olfactory dopamine neurons fire spontaneously at 1-10 Hz and up to 50 Hz or 75 Hz, respectively, in response to somatic current injection [40,50,63,68]. Here, we show that the majority of A11 dopamine neurons exhibited spontaneous rhythmic pacemaking at ∼5 Hz, similar to pacemaking in midbrain and olfactory dopamine neurons. While I_h_ is associated with pacemaking in many populations, including midbrain dopamine neurons [69]; here we report no evidence of I_h_ in A11 dopamine neurons, similar to A12 and olfactory dopamine neurons [40,58,61,63].

In midbrain and olfactory bulb dopamine neurons, the net current at subthreshold potentials which drives pacemaking is often quite small and carried by persistent sodium or calcium influx [63,70,71]. Here, we show that at −50 mV, there was a net inward current of ∼-60 pA carried by sodium influx in pacemaking A11 dopamine neurons, which likely provides the subthreshold drive to fire spontaneously. The quiescent neurons, but not pacemaking neurons, had an outward leak potassium current that outweighed inward current. Whether quiescent neurons had diminished sodium conductance relative to pacemaking neurons remains to be determined.

In neuronal circuits, pacemaking is associated with the maintenance of circadian rhythm [72–74] and the establishment and refinement of sensory and nociceptive circuits [75–77], at least in part due to continual release of modulatory neurotransmitters [78], although many questions remain. In midbrain dopamine neurons, pacemaking is proposed to provide continual dopamine release in axon terminal projections areas. Thus, it is possible that pacemaker activity in A11 dopamine neurons similarly provides steady dopamine input to the spinal cord. Indeed, rodent studies have demonstrated that A11 dopamine neurons are linked to circadian changes in spinal dopamine levels, pain behaviors, and limb movement [10,14], as well as the development of chronic pain [19,52]. How slow pacemaking activity of A11 dopamine neurons translates to spinal dopamine levels and whether there are diurnal changes in ion channel expression or synaptic input remains to be resolved.

### Health Implications

As the only known source of spinal dopamine [3–5], A11 dopamine neurons are implicated in descending pain and itch modulation [3,6–8,19], restless leg syndrome [16,79,80], and motor signaling [11,13]. A11 dopamine neurons also project to the trigeminocervical complex in the brainstem where their activity may contribute to orofacial pain and migraine [20,45,81]. In humans, the mechanisms underlying chronic pain, migraine, and restless legs syndrome remain poorly understood and without adequate treatment options [82,83].

Much of what is known about the functional role of A11 dopamine neurons comes from rodent studies where A11 dopamine neurons are stimulated or destroyed, or inferred by manipulating dopamine receptor signaling in the spinal cord. In healthy animals, electrical stimulation of the A11 region or spinal application of broad dopamine receptor agonists reduces nociceptive transduction and pain behaviors [3,8,90–92]. In rodents, chemically ablating A11 dopamine neurons mimics motor behaviors observed in patients with restless legs syndrome [12,14,87], but can also either increase [9,20,81] or decrease [19,45,52] pain behaviors in experimental models of pain.

In one study, systemic administration of a mu-opioid receptor agonist, morphine, increased the amount of dopamine in the lumbar spinal cord one hour later, suggesting that morphine increases the activity of A11 dopamine neurons [88]. Presynaptic inhibition of GABAergic inputs to A11, and subsequent disinhibition of A11 dopamine neurons was proposed [88]. Indeed, we found that activation of mu-opioid receptors with DAMGO inhibited spontaneous GABAergic synaptic transmission on A11 dopamine neurons. However, activation of mu-opioid receptors also reduced the net inward current driving pacemaking by activating potassium conductance. In cell-attached recording, activation of mu-opioid receptors slowed pacemaking in A11 dopamine neurons, suggesting the net effect of opioids on A11 dopamine neuron activity is inhibitory, but the inhibitory effect desensitized in the continued presence of agonist. *In vivo*, disinhibition may outlast direct activation of potassium current since presynaptic mu-opioid receptors can be resistant to acute desensitization [89]. Opioids can be effective in alleviating pain and symptoms of restless legs syndrome but can also induce itch and hyperalgesia with long-term use. Whether adaptations to A11 dopamine neuron activity contribute to these unwanted effects remains to be tested. Despite evidence that spinal dopamine modulates pain, efforts to target spinal dopamine signaling in human conditions are complicated by the small size of A11 [15] and the difficulty in pharmacologically targeting dopamine receptors without undesirable side effects limiting adherence [90,91].

Lastly, it is well-established that chronic pain conditions are more prevalent and severe in women [92,93]. A prior study demonstrated that male mice have more A11 dopamine neurons and a higher concentration of dopamine in the spinal cord when compared to female mice [94]. In the present study, more A11 dopamine neurons from male mice had pacemaker activity when compared to neurons from female mice, which may facilitate more dopamine release in the spinal cord. Future studies into the synaptic and ion channel mechanisms that govern the action potential firing of A11 dopamine neurons may reveal novel pharmacological targets in the treatment of pain, migraine, and restless legs syndrome.

## Data Availability

The datasets generated during and/or analyzed during the current study are available from the corresponding author on reasonable request.

## Acknowledgements

We would like to thank Dr. Deniz Atasoy (University of Iowa) for providing the TH-IRES-Cre mice (EM:00254) and Lynn Rasmussen for their technical assistance in the retrograde Fluoro-Gold experiments.

## Author Contributions

Conceptualization, A.F.S, K.A.S, S.C.G; Data Curation A.F.S, H.N.R, S.C.G; Formal Analysis A.F.S, H.N.R, S.C.G; Funding Acquisition A.F.S, K.A.S, S.C.G; Investigation A.F.S, H.N.R; Methodology, A.F.S, S.C.G; Project Administration S.C.G; Resources K.A.S, S.C.G; Software, S.C.G; Supervision, K.A.S, S.C.G; Validation, A.F.S, H.N.R, S.C.G; Visualization, A.F.S, H.N.R, S.C.G; Writing – original draft, A.F.S, S.C.G; Writing – review and editing, H.N.R, K.A.S.

## Materials and Methods

### Brain slice preparation and electrophysiological recordings

A11 neurons labeled with tdTomato were visualized on an upright microscope (Olympus, BX51WI) equipped with LED illumination (Cite Xylis, Excelitas). Electrophysiological recordings were made at 35°C with a MultiClamp 700B amplifier (Axon Instruments), Digidata 1550B A/D converter (Axon Instruments) and pClamp 11 software (Molecular Devices). Borosilicate glass electrodes (Warner Instruments) were wrapped with Parafilm and had resistances between 2.1 and 5.8 MΩ when filled with internal solution. Measures of intrinsic and active membrane properties were collected in the absence of antagonists. Loose (5-30 MΩ) cell-attached recordings were made with pipettes filled with modified Krebs buffer and 0.1% Biotin ethylenediamine. Spontaneous excitatory postsynaptic currents were recorded in the presence of picrotoxin (100 μM) to isolate glutamatergic synaptic transmission. GluA/GluK and selectively GluA synaptic transmission was blocked using NBQX (3 µM) or GYKI-52466 (50 µM), respectively [1,2]. Spontaneous inhibitory postsynaptic currents were recorded in the presence of NBQX (3 μM) to isolate GABAergic synaptic transmission, with a high-chloride containing internal solution (calculated E_Cl_: −1.73 mV): (in mM), K-methyl sulfate (14), KCl (111), NaCl (5.3), K-HEPES (7.07), MgCl_2_ (4.06), CaCl_2_ (0.2245), EGTA (0.45), Na-GTP (0.26), Na-ATP (4.87), Na-creatine phosphate (4.59), and 0.1% Biotin ethylenediamine [1,3], pH 7.25 with KOH, mOsm ∼278. GABA_A_ synaptic transmission was blocked using picrotoxin (100 µM). DAMGO was used to activate mu-opioid receptors (10 µM). All drugs were applied by bath application.

### Retrograde tracing

C57BL6/J mice were anesthetized with isoflurane (5% induction, 0.5-5% maintenance, 1.5 L/min). A laminectomy was performed to expose the lumbosacral spinal cord and a piece of gel foam soaked in 4% Fluoro-Gold (Fluorochrome) was applied to the dorsal aspect of the spinal cord at the L4-L6 level. Incisions were closed and the mice recovered for 3 weeks. Mice were euthanized and perfused transcardially with heparinized saline followed by ice-cold 4% paraformaldehyde (PFA) in phosphate buffer (PB, pH 7.4). Brains and spinal cords were extracted and post-fixed overnight or 48 h, respectively, in 4% PFA. Tissue was cryoprotected in increasing solutions of sucrose in PBS (10%, 20%, 30%) and then frozen in optimal cutting temperature (OCT) compound and sliced coronally in 40-µm sections and processed for immunohistochemistry. To verify Fluoro-Gold application and location in the lumbosacral spinal cord, coronal 30 µm slices were made on a cryostat, directly mounted onto slides, and immunostained for Fluoro-Gold as described below. Spinal cord sections were inspected to verify several millimeters of Fluoro-Gold at the level of the lumbosacral spinal cord but not the thoracic spinal cord.

### Immunohistochemistry and confocal microscopy

#### For enzyme expression in A11

C57BL/6J mice were euthanized and perfused transcardially with 5% sucrose in diH_2_O followed by ice-cold 4% paraformaldehyde (PFA) in phosphate-buffered saline (PBS, pH 7.4). Brains were post-fixed overnight in 4% PFA then sliced coronally in 60-μm sections. Free-floating slices were permeabilized and blocked in PBS with 0.2% Triton-X, 5% normal donkey serum, and 5% BSA for 1 h and 45 min. Slices were incubated overnight in primary antibody; chicken anti-tyrosine hydroxylase (TH) (1:500, Aves labs TYH) or goat anti-tryptophan hydroxylase 2 (TpH2, 1:1000, Abcam AB121013), in 0.2% Trion-X and 1% donkey serum. Slices were washed then incubated in donkey anti-chicken Alexa Fluor 594 (1:500, Jackson Labs 703-585-155) or anti-goat Alexa Fluor 697 (1:1000, Jackson Labs 705-585-003) for 2 h. Following washing, slices were incubated in primary antibody; rabbit anti-amino acid decarboxylase primary (AADC, 1:500, Novus Biologicals ab9702) or rabbit anti-dopamine-β-hydroxylase (DβH, 1:3000, Immunostar 22806), 1% donkey serum, and 5% BSA for 44 hours. Slices were washed then incubated in donkey anti-rabbit Alexa Fluor 488 (1:500, Jackson Labs 711-545-152) for 2 h. Lastly, slices were washed, mounted, and cover-slipped with Flouromount-G™ with DAPI. Fluorescent images (10× or 40× magnification) were obtained with an Axiovert 100 confocal microscope (Zeiss) 1-6 images per section were taken on 3-9 sections per animal. 1.08-1.23 µm thick z-planes were taken in sufficient number to capture the full thickness of the tissue (6-36 z-planes/image)

#### For retrograde tracing

Slices with Fluoro-Gold retrograde tracing were prepared as described above and permeabilized in 0.05% Triton X-100 for 5 min, then blocked in 5% normal goat serum for 30 min. Slices were incubated overnight in chicken anti-TH (1:1000, Millipore Sigma, Ab9702) and rabbit anti-Fluoro-Gold (1:500, Fluorochrome) in 0.05% Triton-X-100 and 1% goat serum. Following washing, slices were incubated in goat anti-chicken Alexa Fluor 647 (1:1000, ThermoFisher A32933) and goat anti-rabbit biotin (1:1000, Jackson Immunno 111-066-144) for 2 h. Slices were washed then incubated in streptavidin 405 (1:1000, Invitrogen 2539805) for 2 h. Lastly, slices were washed, mounted, and cover-slipped with Flouromount-G™. Fluorescent images (40× magnification) were obtained with an Axiovert 100 confocal microscope (Zeiss). Two images per section were taken on four sections per animal. 1.12 µm thick z-planes were taken in sufficient number to capture the full thickness of the tissue (13-32 z-planes/image)

#### For Cre line validation

TH-Cre-1::Ai14, TH-IRES-Cre::Ai14, or TH-2A-CreER::Ai14 mice were euthanized and perfused transcardially with heparinized saline followed by ice-cold 4% paraformaldehyde (PFA) in phosphate buffer (PB, pH 7.4). Brains were extracted and post-fixed overnight in 4% PFA. Tissue was cryoprotected in increasing solutions of sucrose in PBS (10%, 20%, 30%) and then frozen in optimal cutting temperature (OCT) compound and sliced coronally in 40 µm sections. Sections containing A11 were permeabilized in 0.05% Triton X-100 for 5 min, then blocked in 5% normal goat serum for 30 min. Slices were incubated overnight in chicken anti-TH (1:1000, Millipore Sigma, Ab9702) in 0.05% Triton-X-100 and 1% goat serum. Following washing, slices were incubated in goat anti-chicken Alexa Fluor 488 (1:1000, ThermoFisher A32931) for 2 h. Lastly, slices were washed, mounted, and cover-slipped with Flouromount-G™ with DAPI. Fluorescent images (28× magnification) were obtained with an Axiovert 100 confocal microscope (Zeiss). Four images were taken on three sections per animal. Four 1 µm thick z-planes were taken of the top and bottom 4 µm of each section to avoid false negatives (a tdTomato^+^ neuron appearing TH^-^ due to incomplete antibody penetrance in deep tissue).

#### For electrophysiology

Slices used for electrophysiological recordings were placed in 4% PFA for 1 h following recordings. Slices were washed, then permeabilized and blocked in 0.5% Triton-X, 10% NGS in PBS for 5 h. Slices were incubated in rabbit-anti TH (1:1000, Millipore Sigma, AB152) antibody overnight. Following washing, slices were incubated in goat anti-rabbit Alexa Fluor 488 (1:1000, Thermofisher A32731) for 2 h, washed, then incubated in Streptavidin Alexa-Fluor 647 (1:1000, Thermofisher, S21374) for 2 h. Slices were washed, mounted, and cover-slipped with Flouromount-G™ with DAPI. Fluorescent images (40× magnification) were obtained with an Axiovert 100 confocal microscope (Zeiss). For whole-cell recordings, the recorded neuron was identified by labeling with Alexa-Flour 647. For cell-attached recordings, due to poor biotin filling from loose seal recordings [4], fluorescent images of recorded neurons were acquired live with a 40× water-dipping objective, then fixed as above for post-hoc validation of TH immunostaining. The recorded neuron was identified based on location in A11 and the morphology of the recorded and nearby neurons. 1.88 µm thick z-planes were taken of the recorded neuron in sufficient number to capture all visible dendrites (8-75 z-planes/image).

#### Immunohistochemistry Image Analysis

Immunostaining of TH, AADC, TpH2, DβH, FG, and tdTomato were counted using FIJI (ImageJ) software. Specificity and penetrance were measured as the percentage of tdTomato^+^ neurons expressing TH and the percentage of TH^+^ neurons expressing tdTomato, respectively. Morphology measurements were calculated with the Simple Neurite Tracer (SNT) v4.2.1 in the Neuroanatomy plug-in for FIJI. Primary and secondary dendrites were manually identified and labeled using SNT with snapping and A* search algorithm enabled. Dendritic arbor length measurements, counts, and 3D reconstructions were made using SNT’s path properties and reconstruction features.

## Materials

MK-801 and GYKI 52466 were obtained from Tocris. Picrotoxin and DAMGO were from ThermoFisher. NBQX was obtained from Cayman Chemicals. All other reagents were obtained from Sigma Aldrich.

## Data and statistical analyses

Data were analyzed using Clampfit 11.1 and Igor 6 (Wavemetrics) using DataAccess (Bruxton). Using Clampfit 11.1, cell capacitance was calculated by fitting the capacitive transient (ι1V=5 mV) with a double exponential and then using the resulting ι−-fast and access resistance for the same steps. Neurons were determined to be pacemakers if they sustained firing at a rate ≥1 Hz. Most quiescent neurons never spontaneously fired (n=15), however neurons that spontaneously fired only a few action potentials at <1 Hz (n=11) were classified as quiescent. Spontaneous EPSCs were detected using the semi-automated template search in Clampfit with a template match threshold of 5.5, using two templates generated from a single ‘fast’ EPSC with a ι−-decay of 1.72 ms and a ‘slow’ EPSC with a ι−-decay of 4.45 ms. Spontaneous IPSCs were similarly detected using a template generated by a single IPSC with a ι−-decay of 6.28 ms. Detected events were manually checked. Time to peak and !-decay were determined by averaging all fast and slow events for each cell and measuring time from beginning of event to peak or fitting a single exponential on the average trace, respectively. Only neurons with >10 fast and >10 slow sEPSCs were included in the ratio analysis. No difference in sEPSC frequency, amplitude, or kinetics was observed when compared by firing type or sex, so data were pooled. Electrophysiology data are presented as representative traces, in scatter plots where each point is an individual cell, or line graphs with mean±SEM. n=number of cells as biological replicates unless otherwise stated. Data sets with n>30 were tested for normality with a Shapiro-Wilk test, otherwise non-parametric statistical tests were performed as sample size was too small to evaluate normality. Paired tests were used for within-cell comparisons, whereas between-cell comparisons were made with unpaired tests or ordinary two-way ANOVAs with Sidak multiple comparisons where appropriate were used. When testing for correlation between two measures, a simple linear regression was performed. A difference of p<0.05 was considered significant. Statistical analyses were performed using GraphPad Prism (GraphPad Software).

Means±SEM, n=82&59 total neurons (pacemaker, quiescent, or depolarization blocked) from 22 male mice and 24 female mice, respectively. For pacemaking neurons comparisons, males: n=74-75; Females: n=35-38. Chi squared test (% pacemaking), unpaired t-test with Welch’s correction (threshold and AHP), and non-parametric Mann-Whitney tests (frequency, peak height, and half-width). Bold-type designates statistical significance.

---

**Supplementary Table S1.** Characteristics of pacemaking A11 dopamine neurons from male and female mice.

| Sex | Males | Females | p-value |
| --- | --- | --- | --- |
| pacemaking (%) | 91.5 | 61.0 | **<0.0001** |
| spontaneous frequency (Hz) | 5.5±0.4 | 5.4±0.5 | 0.9301 |
| Threshold (mV) | -39.8±0.5 | -39.5±0.5 | 0.6436 |
| Peak height (mV) | 80.7±1.2 | 80.5±1.5 | 0.6627 |
| Half-width (ms) | 1.3±0.04 | 1.4±0.09 | 0.1430 |
| AHP (mV) | -65.7±0.8 | -65.7±0.8 | 0.9967 |

**Supplementary Fig S1.**
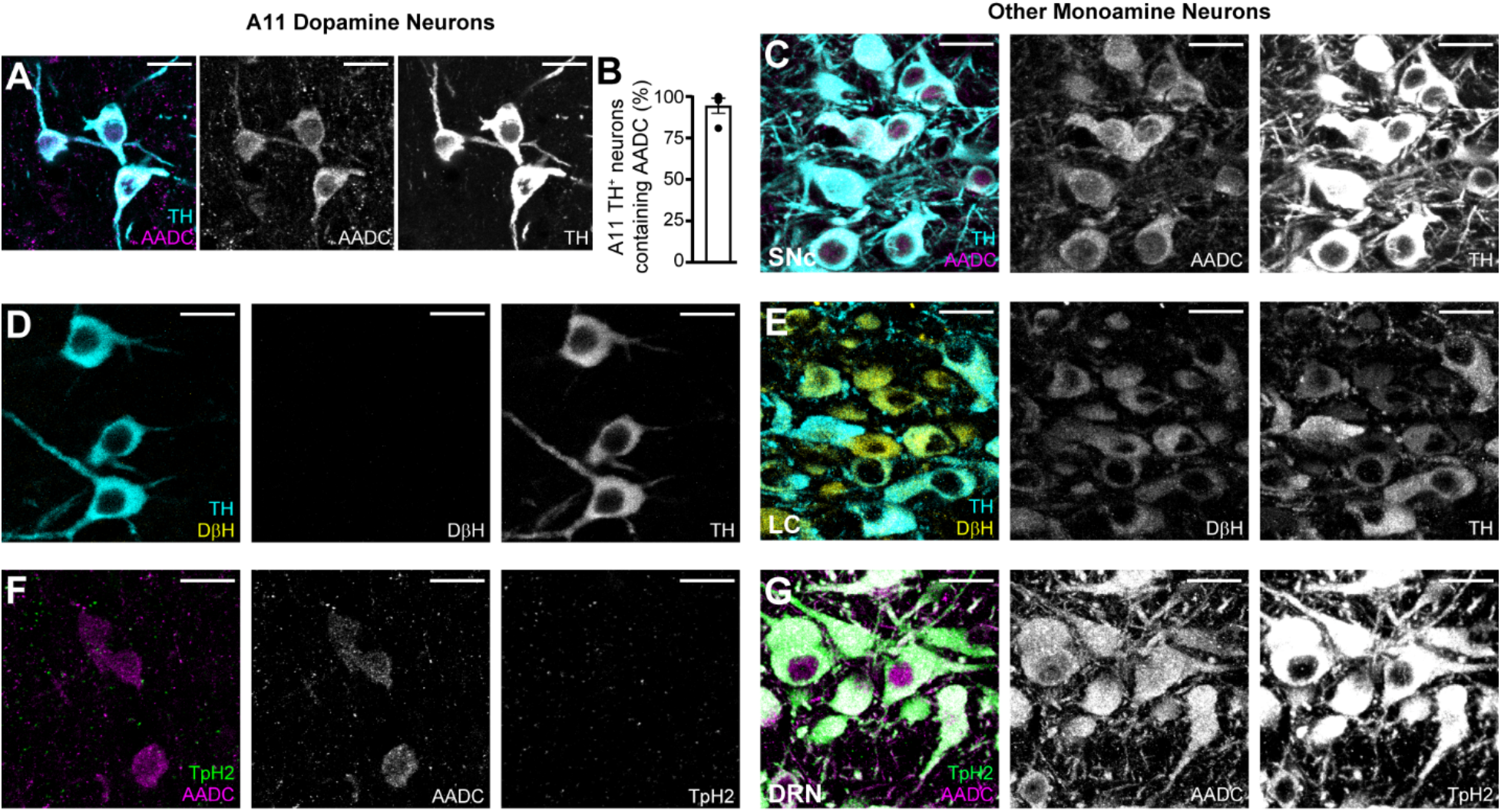
A11 neurons do not synthesize noradrenaline or serotonin. **A** Representative maximum intensity projection confocal image of TH^+^ neurons in A11. Sections were immunostained for tyrosine hydroxylase (TH, cyan) and L-amino acid decarboxylase (AADC, magenta) demonstrating expression of AADC in TH^+^ neurons. **B** Percentage of TH^+^ neurons that contained L-amino acid decarboxylase (AADC) in A11. n=4 mice, 362 TH^+^ neurons. **C** Representative maximum intensity projection confocal image of TH^+/^AADC^+^ neurons in the substantia nigra pars compacta (SNc) as a positive control. **D** Representative maximum intensity projection confocal image of TH^+^ neurons in A11. Sections were immunostained for tyrosine hydroxylase (TH, cyan) and dopamine-β-hydroxylase (DβH, yellow) demonstrating no expression of DβH in TH^+^ neurons. n=4 mice, 288 TH^+^ neurons. **E** Representative maximum intensity projection confocal image of TH^+^/DβH^+^ neurons in the locus coeruleus (LC) as a positive control. **F** Representative maximum intensity projection confocal image of AADC^+^ neurons in A11. Sections were immunostained for L-amino acid decarboxylase (AADC, magenta) and tryptophan hydroxylase 2 (TpH2, green) demonstrating no expression of TpH2 in AADC^+^ neurons. n=4 mice, 293 AADC^+^ neurons. **G** Representative maximum intensity projection confocal image of AADC^+^/TpH2^+^ neurons in the dorsal raphe nucleus (DRN) as a positive control. Scale bars 20 μm. Line and error bars represent Means±SEM.

**Supplementary Fig S2.**
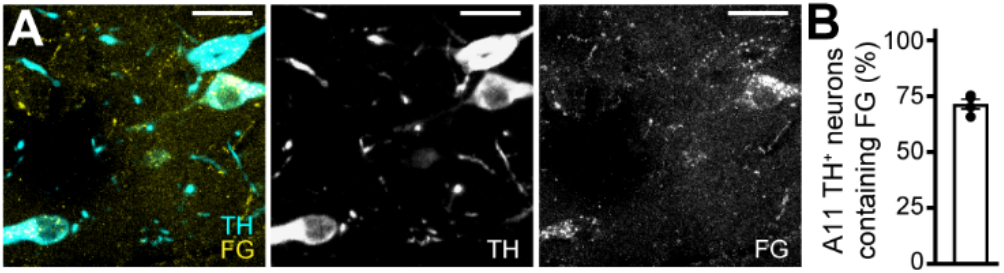
A11 dopamine neurons project to the spinal cord. **A** Representative maximum intensity projection confocal image of TH^+^ neurons in A11. Three weeks after Fluoro-Gold was placed at the lumbosacral level of the spinal cord, brain sections containing A11 were immunostained for TH (cyan) and Fluoro-Gold (FG, yellow) labeling TH^+^ neurons that project to the lumbosacral section of the spinal cord; scale bars 20 μm. **B** Percentage of TH^+^ neurons that contained Fluoro-Gold (FG) in A11. n=4 mice, 337 TH^+^ neurons. Line and error bars represent Means±SEM.

**Supplementary Fig S3.**
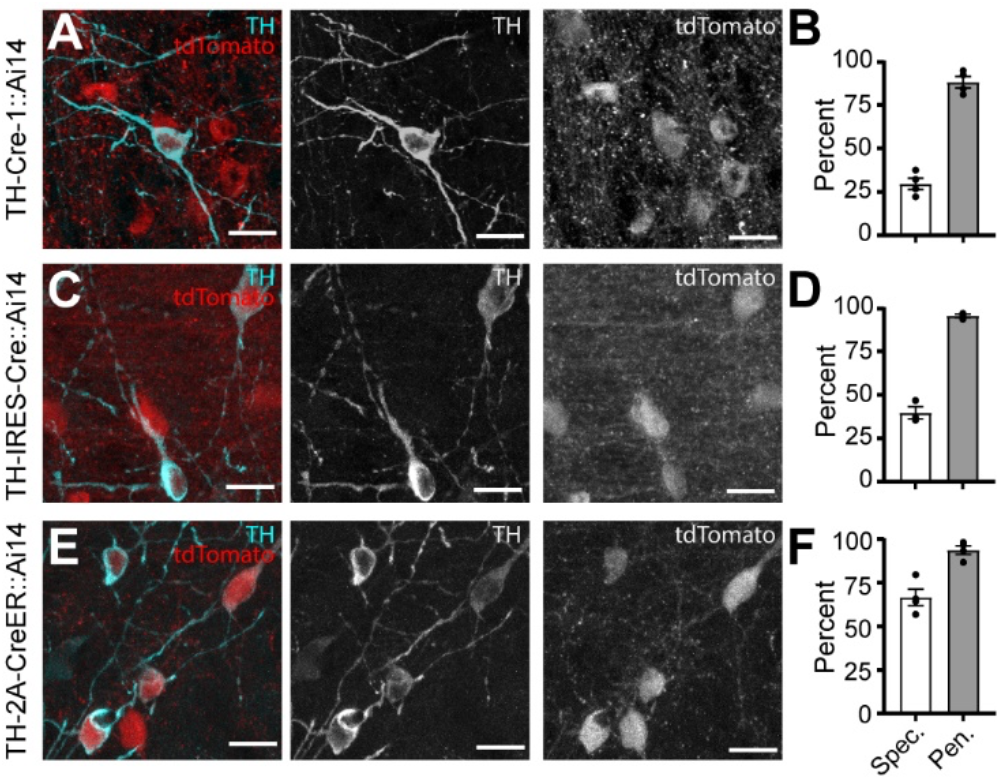
Available Cre lines have variable specificity A11 dopamine neurons. **A** Maximum intensity projection confocal image of dopamine neurons in A11 in TH-Cre-1::Ai14 mice, which produced cytosolic tdTomato expression in Cre^+^ neurons (red). Slices were immunostained for TH (cyan). Scale bars 20 μm. **B** Specificity (percent of tdTomato^+^ neurons colocalizing with TH) and penetrance (percent of TH^+^ neurons colocalizing with tdTomato) in A11 in TH-Cre-1::Ai14 mice. n=4 mice, 586 tdTomato^+^ neurons, 197 TH^+^ neurons. **C** Maximum intensity projection confocal image of dopamine neurons in A11 in TH-IRES-Cre::Ai14 mice (red). Slices were immunostained for TH (cyan). Scale bars 20 μm. **D** Specificity and penetrance in A11 in TH-IRES-Cre::Ai14 mice. n=3 mice, 303 tdTomato^+^ neurons, 125 TH^+^ neurons. **E** Maximum intensity projection confocal image of dopamine neurons in A11 in TH-2A-CreER::Ai14 mice (red). Slices were immunostained for TH (cyan). Scale bars 20 μm. **F** Specificity and penetrance in A11 in TH-2A-CreER::Ai14 mice. n=4 mice, 243 tdTomato^+^ neurons, 169 TH^+^ neurons. Line and error bars represent Means±SEM.

**Supplementary Fig S4.**
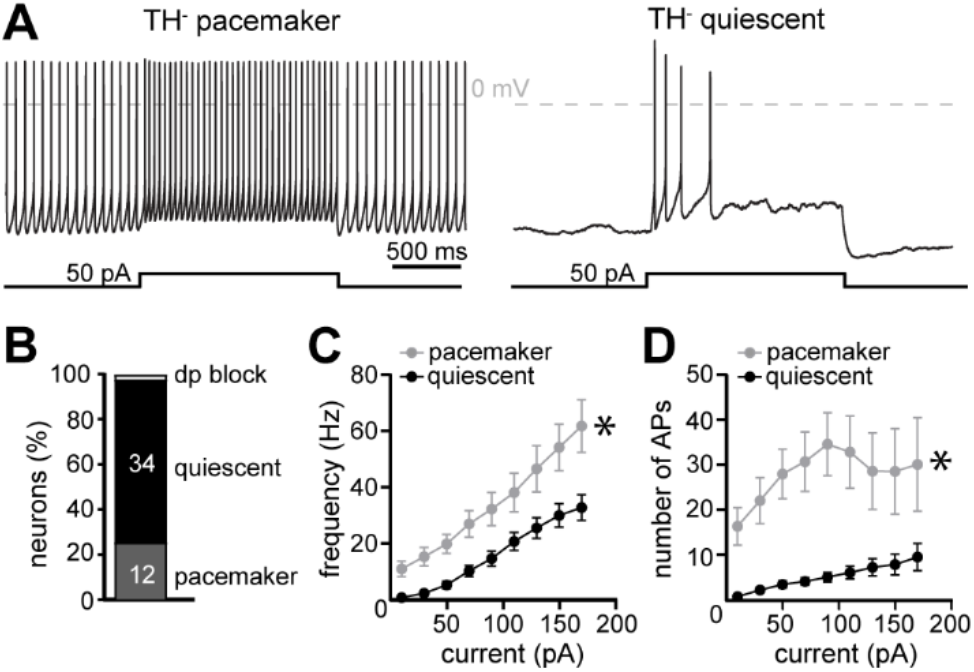
A11 TH^-^ neurons also can fire action potentials in a rhythmic pacemaker manner. **A** Representative current-clamp traces of A11 TH^-^ neurons that fired action potentials spontaneously (‘pacemakers’, left) or were silent (‘quiescent’, right). Both types fired action potentials during somatic current injection (50 pA, 1.5 s). **B** Percentage of A11 TH^-^ neurons that fired spontaneously (pacemaker), were silent (quiescent), or in depolarization block (dp). Inset numbers represent n of neurons. n=47. **C** Pacemaker neurons fired at a higher frequency during a somatic current injection than quiescent neurons. (p<0.0001, n=12&34). **D** Number of action potentials was increased significantly in pacemaker neurons. (p<0.0001, n=12&34). Line and error bars represent means±SEM. Data were analyzed by ordinary two-way ANOVA, main effect of type. * denotes statistical significance.

**Supplementary Fig S5.**
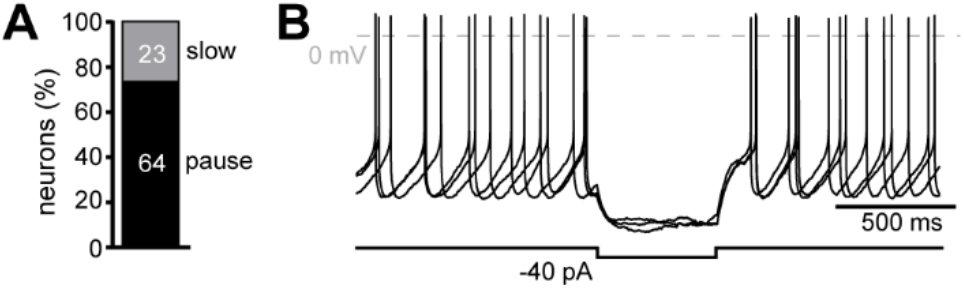
Prolonged hyperpolarization inhibits pacemaking in most A11 dopamine neurons. **A** Percentage of pacemaking A11 dopamine neurons that paused or slowed firing in response to negative current injection (−40 pA). Inset numbers represent n of neurons (n=87). **B** Representative current-clamp traces of a pacemaking A11 dopamine neurons that paused firing in response to negative current injection (−40 pA, 0.5 s).

**Supplementary Fig S6.**
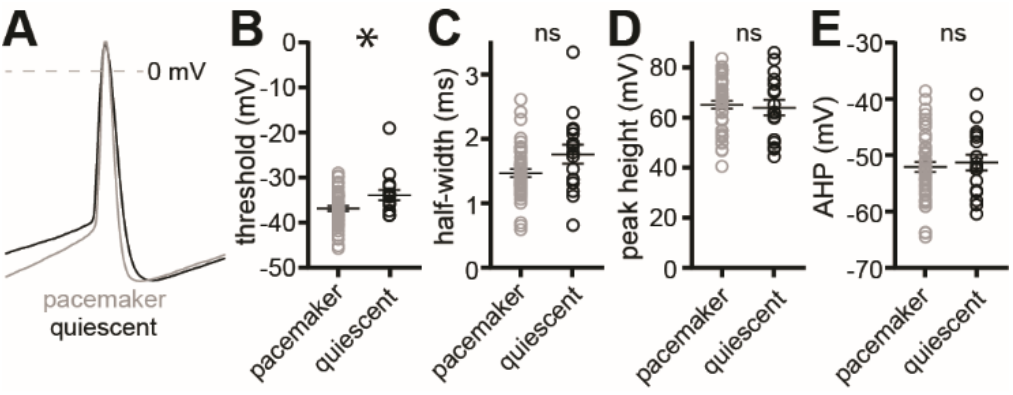
Evoked action potential waveform parameters of pacemaker and quiescent A11 dopamine neurons. **A** Representative action potentials induced by somatic current injection (150 pA) from a pacemaker (grey) and quiescent (black) A11 dopamine neuron aligned at peaks. **B** Quiescent neuron action potentials had a more depolarized threshold than pacemaker neurons (p=0.0349) **C** The half-width (measured at mid-point between peak and trough) did not differ between pacemaker and quiescent neurons (p=0.0557). **D** The peak height of the action potential did not differ between pacemaker and quiescent neurons (p=0.7052). **E** The magnitude of the after-hyperpolarization (AHP) did not differ between pacemaker and quiescent neurons (p=0.7175). All panels n=44 pacemaking and 17 quiescent neurons, 31 mice. All measures were made with the 2^nd^ action potential. Data were analyzed with a non-parametric unpaired t-test (B-E). Line and error bars represent means±SEM. * denotes statistical significance, ‘ns’ indicates not significant.

**Supplementary Fig S7.**
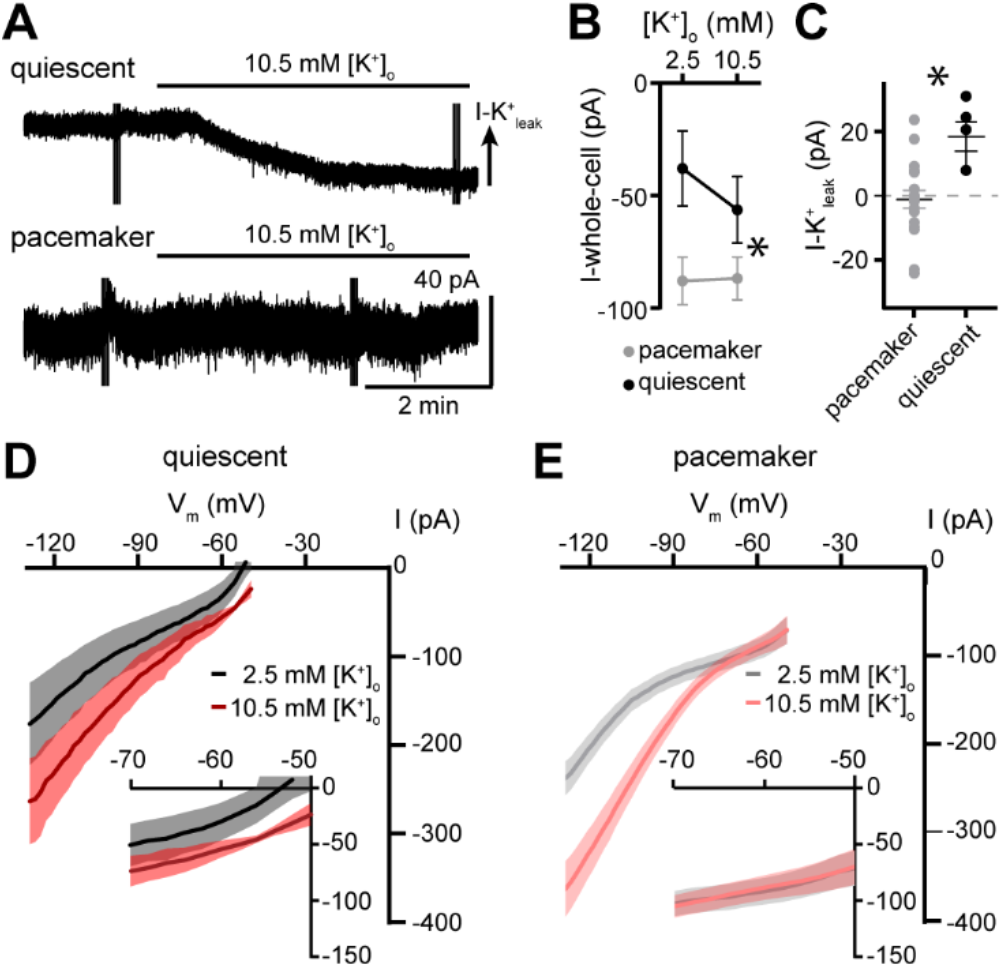
Quiescent A11 dopamine neurons have a potassium leak conductance at subthreshold potentials. **A, B** Increasing extracellular potassium to 10.5 mM produces an apparent inward current due to a reduction in leak potassium current in quiescent but not pacemaker neurons, shown in **A,** representative whole-cell voltage-clamp recordings (voltage ramps truncated for clarity) and **B**, grouped data (Two-way repeated measures ANOVA, interaction, p=0.0033, n=5&19). **C** Plot of the magnitude of the leak potassium current measured at V_hold_ −65 mV in pacemaker and quiescent neurons (Mann-Whitney test, p=0.0020, n=5&19). **D, E** Current-voltage plots generated by a voltage-ramp from −50 mV to −130 mV (0.2 mV/ms) in control conditions (2.5 mM [K^+^]_o_) and in 10.5 mM [K^+^]_o_, showing a reduction in leak potassium current at in (**D**) quiescent (n=5) but not (**E**) pacemaker neurons (n=19). Line and error bars represent means±SEM. * denotes statistical significance.

**Supplementary Fig S8.**
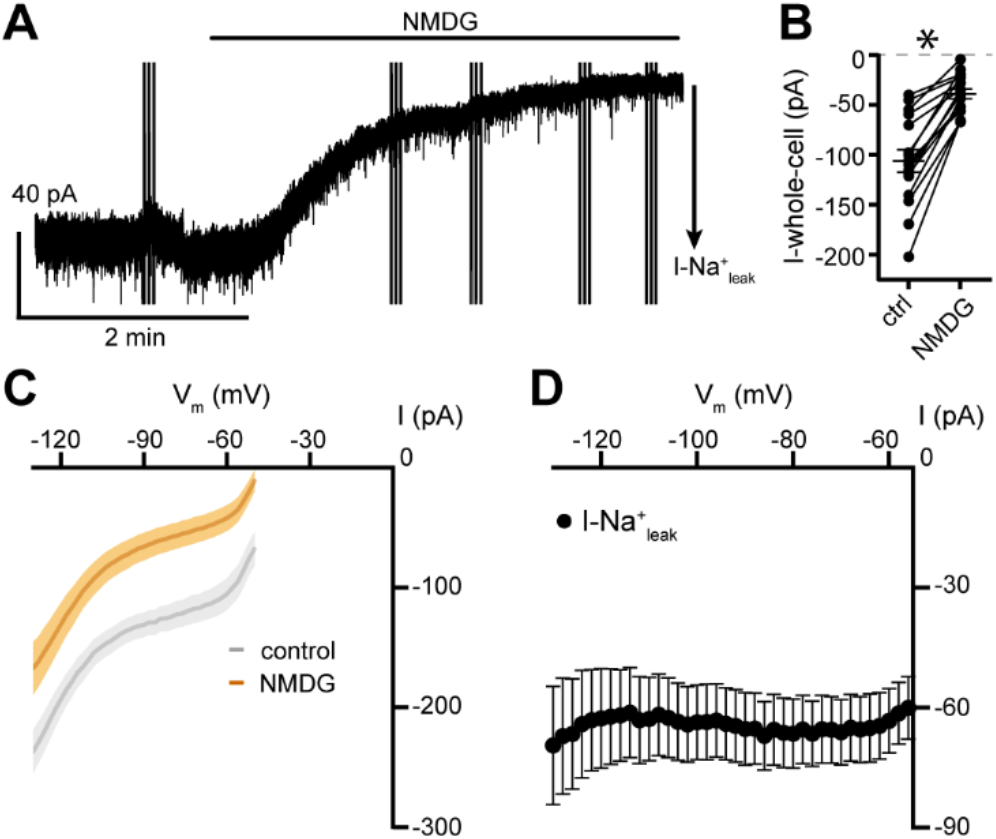
Subthreshold inward current in pacemaking A11 dopamine neurons is carried primarily by sodium entry. **A, B** Replacing 115 mM extracellular sodium with NMDG produces an apparent outward current due to a reduction in leak sodium current in pacemaker neurons, shown in **A,** representative whole-cell voltage-clamp recording (voltage ramps truncated for clarity) and **B**, grouped data (Wilcoxon test, p<0.0001, n=16). **C, D** Current-voltage plots generated by a voltage-ramp from −50 mV to −130 mV (0.2 mV/ms) in (**C**) control conditions and NMDG, showing a reduction in leak sodium current (I-Na^+^_leak_) in pacemaker neurons (n=16), also illustrated in (**D**) which indicates little to no voltage-dependence in the current. Line and error bars represent means±SEM. * denotes statistical significance.

**Supplementary Fig S9.**
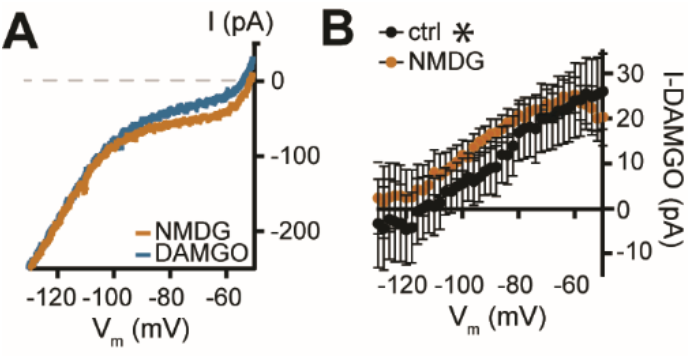
Removal of sodium has little effect on mu-opioid receptor-suppression of net inward current. **A** Example current-voltage plots generated by a voltage-ramp from −50 mV to −130 mV (0.2 mV/ms) in NMDG and after application of a mu-opioid receptor agonist, DAMGO (10 μM) in NMDG in a single neuron. **B** Current-voltage plots of the DAMGO-induced current (I-DAMGO) in control conditions (n=8, sodium-containing) and in NMDG (n=7, Ordinary two-way ANOVA, main effect of sodium concentration, p<0.0041). Line and error bars represent means±SEM. * denotes statistical significance, ‘ns’ indicates not significant.

**Supplementary Fig S10.**
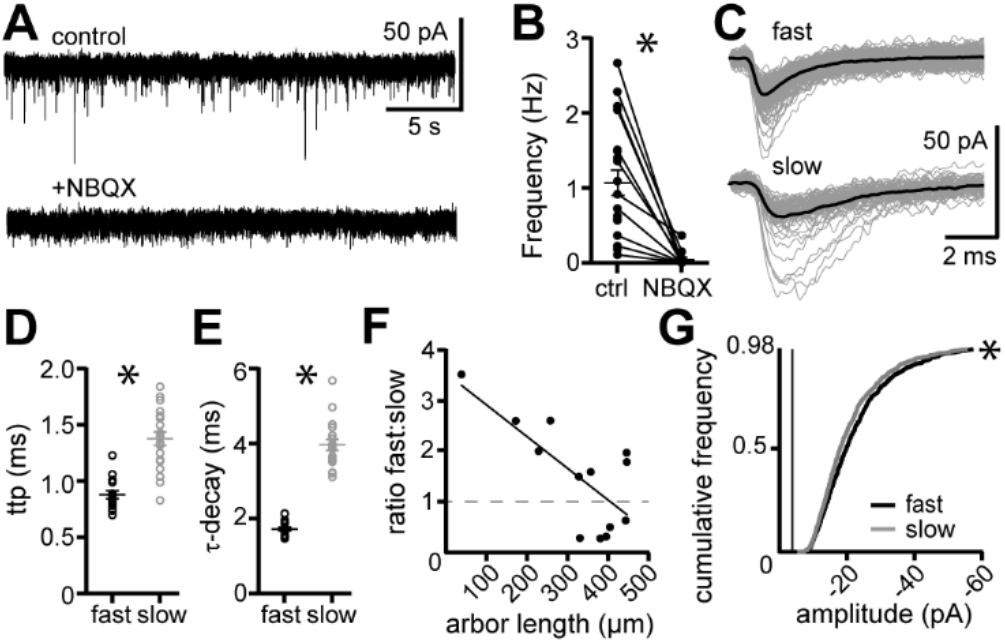
A11 dopamine neurons receive ionotropic glutamatergic input. **A** Representative trace of spontaneous EPSCs (sEPSCs) in an A11 dopamine neuron in control conditions and after application of NBQX (3 μM). **B** Frequency of sEPSCs; there was a significant reduction in sEPSCs following application of NBQX (3 μM, Wilcoxon matched-pairs signed-rank test, p=0.001, n=11-20 neurons, 7-13 mice). **C** Representative sEPSCs from an A11 dopamine neuron. Two types of sEPSCs were detected; fast and slow. **D** Fast sEPSCs had a significantly shorter time-to-peak (ttp) when compared to slow sEPSCs (Mann-Whitney test, p<0.0001, n=20 neurons, 13 mice). **E** Fast sEPSCs had a significantly shorter τ-decay when compared to slow sEPSCs (Mann-Whitney test, p<0.0001, n=20, 13 mice). **F** Ratio of fast to slow sEPSCs was correlated to dendritic arbor length (simple linear regression, R^2^=0.5292, p=0.0048, n=13 neurons). Dashed line represents the unity line where the number of fast sEPSCs equals the number of slow sEPSCs. **G** Cumulative frequency of amplitude for fast and slow sEPSCs. Box indicates mean noise as the lowest limit of detection (−4 pA). The y-axis is truncated to show 98% of events (Kolmogorov-Smirnov test, p=0.0002, n=1267-1325 events from 20 neurons and 13 mice). Line and error bars represent means±SEM. * denotes statistical significance.

**Supplementary Fig S11.**
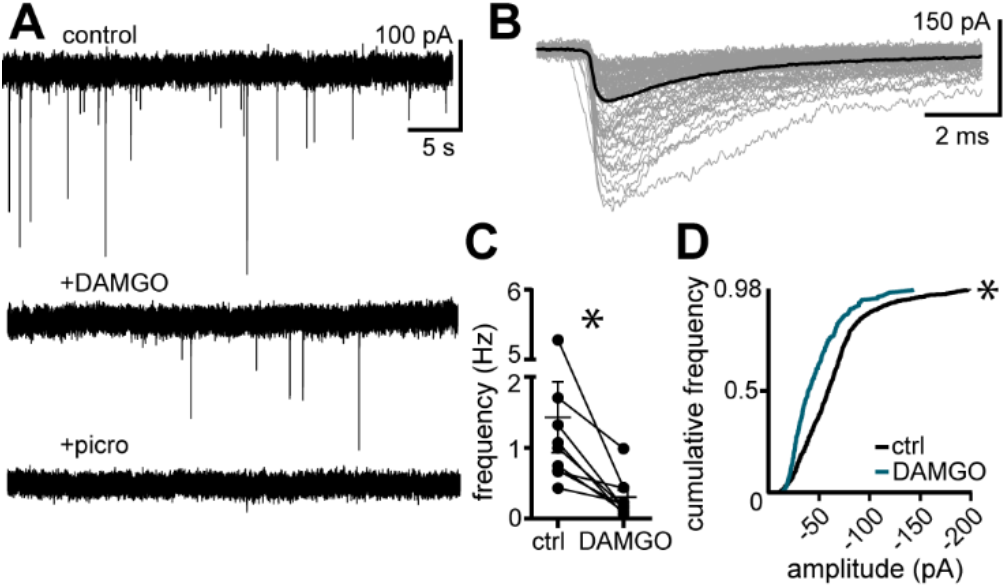
Activation of mu-opioid receptors inhibits GABA_A_ synaptic input onto A11 dopamine neurons. **A** Representative trace of spontaneous inhibitory postsynaptic currents (sIPSCs) recorded using a high-chloride internal solution in an A11 dopamine neuron in control conditions, after application of DAMGO (10 μM), and after application of picrotoxin (100 μM). **B** Representative traces of sIPSCs recorded from a single neuron in control conditions. **C** DAMGO produced a significant reduction in sIPSCs (Wilcoxon matched-pairs signed-rank test, p=0.0039, n=9 neurons from 5 mice). **D** DAMGO produced a significant reduction in the amplitude of sIPSCs (Kolmogorov-Smirnov test, p<0.0001, n=233-1337 sIPSCs from 9 neurons and 5 mice, the y-axis is truncated to show 98% of sIPSCs). Line and error bars represent means±SEM, * denotes statistical significance.

